# Quantifying *in vivo* collagen reorganization during immunotherapy in murine melanoma with second harmonic generation imaging

**DOI:** 10.1101/2023.11.09.566407

**Authors:** Alexa R. Heaton, Nathaniel J. Burkard, Paul M. Sondel, Melissa C. Skala

## Abstract

**Significance:** Increased collagen linearization and deposition during tumorigenesis can impede immune cell infiltration and lead to tumor metastasis. Although melanoma is well studied in immunotherapy research, studies that quantify collagen changes during melanoma progression and treatment are lacking.

**Aim:** Image *in vivo* collagen in preclinical melanoma models during immunotherapy and quantify the collagen phenotype in treated and control mice.

**Approach:** Second harmonic generation imaging of collagen was performed in mouse melanoma tumors *in vivo* over a treatment time-course. Animals were treated with a curative radiation and immunotherapy combination. Collagen morphology was quantified over time at an image and single fiber level using CurveAlign and CT-FIRE software.

**Results:** In immunotherapy-treated mice, collagen reorganized toward a healthy phenotype, including shorter, wider, curlier collagen fibers, with modestly higher collagen density. Temporally, collagen fiber straightness and length changed late in treatment (Day 9 and 12) while width and density changed early (Day 6) compared to control mice. Single fiber level collagen analysis was most sensitive to the changes between treatment groups compared to image level analysis.

**Conclusions:** Quantitative second harmonic generation imaging can provide insight into collagen dynamics *in vivo* during immunotherapy, with key implications in improving immunotherapy response in melanoma and other cancers.

## 1 Introduction

The extracellular matrix (ECM) is a major component of the tissue microenvironment, providing a scaffold for surrounding cells and regulating cell behavior. Within the extracellular matrix, collagen is a structural protein that comprises 30% of the total protein mass in the human body and is involved in several key biological processes.^1^ In normal tissue, the ECM constantly remodels and repairs, synthesizing new collagen proteins to replace the older, degraded collagen. This process is highly regulated by a precise balance of metalloproteinases (MMPs) and MMP inhibitors.^2,3^ Within the context of cancer, homeostasis is dysfunctional and cancer cells secrete excess amounts of MMPs, degrading the basement membrane and promoting malignant cell invasion into the interstitial matrix.^1,3^ As the tumor progresses, cancer associated fibroblasts (CAFs) secrete excess type I and type II fibrillar collagen, and remodel the collagen morphology within the ECM.^4^ Often, ECM morphology reconstruction results in linearized collagen and increased stiffening within cancer tissue.^1,2^ These phenomena have been categorized as tumor associated collagen signatures (TACS) that summarize the collagen changes that often occur during tumorigenesis: increased deposition or density (TACS-1), more “taut” or straight fibers (TACS-2), and increased fiber alignment (TACS-3).^5,6^ These morphological changes within the tumor microenvironment affect tumor cell migration and metastasis out of the tumor while impacting immune cell recruitment and infiltration into the tumor.^1,2,5,7^

Melanoma is the deadliest of all skin cancers and accounted for 325,000 new cases with an 18% fatality rate globally in 2020.^8,9^ Current predictions expect diagnoses (+50%) and fatalities (+68%) to continue to increase over the next 17 years.^8^ This cancer type affects a wide population with metastatic disease usually driving poor outcomes. With the emergence of somewhat effective immune checkpoint blockade immunotherapy, advanced melanoma prognosis has dramatically improved over the last 25 years, from 9-month median survival rates to a substantial fraction of patients achieving durable cures.^10,11^ Immune checkpoint inhibitors have exhibited especially potent results against melanoma with the best outcomes from combination therapy versus monotherapy.^11,12^ Here, we pursue one such combination strategy including external beam radiation therapy, immune checkpoint inhibitor, and a novel immunocytokine. We have previously shown that this triple combination can cure large GD2+ melanomas in the majority of treated mice.^13–16^ Here, we continue our investigation of this murine B78 melanoma model and expand our inquiry into the tumor microenvironment with a focus on collagen.

Second harmonic generation (SHG) imaging can be used to visualize collagen in its endogenous, label-free state. SHG is a nonlinear optical scattering phenomenon that occurs when two identical photons scatter off a noncentrosymmetric material, collagen here, producing a single photon with exactly twice the energy of the initial photons. As a result of this frequency doubling, SHG signals are always generated at half the excitation wavelength.^17–19^ SHG imaging has emerged as a valuable method to image collagen *in vitro* and *in vivo* due to its high contrast and specificity.^20–24^ One way to quantify collagen morphology from these images is through software developed by the Eliceiri lab at the University of Wisconsin: CurveAlign and CT-FIRE.^6,8,25–27^ These software packages input raw SHG images and quantify several different parameters of collagen morphology including coefficient of alignment, density, straightness, length, and width. In addition to the CurveAlign and CT-FIRE parameters, collagen morphology also varies macroscopically between straight and curly fibers, which can be qualitatively scored for each field of view (FOV).

Although there is significant activity in melanoma immunotherapy research, we are not aware of any studies that have quantified collagen changes during melanoma treatment with immunotherapy, despite collagen comprising 70% of the skin.^28^ In addition, melanoma has been shown to be incredibly fibroblast rich, with human melanoma tumors recruiting activated CAFs.^29–32^ As CAFs are one of the key cell types implicated in pro-tumor collagen and MMP deposition, we anticipate collagen changes within melanoma may be informative to patient prognosis and survival.^29,32–34^ Prior work has used SHG imaging to probe collagen changes primarily in breast ^6,35–42^ and pancreatic cancer^43–48^ with some analyses in melanoma.^22,49,50^ We aim to expand this analysis to melanoma in the context of radiotherapy and immunotherapy treatment.

Here, we quantify collagen morphology features from *in vivo* SHG images of mouse melanoma tumors during combination radiation and immunotherapy versus treatment with phosphate buffered saline (PBS) vehicle. We examine collagen reorganization and phenotypic changes with temporal context and use the quantitative features from CurveAlign and CT-FIRE software to evaluate collagen changes at the field of view level and the single fiber level.

## 2 Methods

### 2.1 Mouse Model

#### 2.1.1 Preparation of Mouse Tumor Model

Animals were housed and treated under an animal protocol approved by the Institutional Animal Care and Use Committee at the University of Wisconsin, Madison. Genetically modified C57BL/6 mice (Jax 000664), with an mCherry reporter within their CD8 T cells, were created at the University of Wisconsin Genome Editing and Animal Models core and used for these studies. The data presented here will not discuss the mCherry expressing CD8 T cells as the focus is just on collagen changes. A separate manuscript, in preparation, will feature the T cell data obtained from these radio-immunotherapy treated mice. Mice were successfully bred and maintained by the University of Wisconsin Biomedical Research Model Services. Male and female mice were used in all studies.

B78-D14 (B78) melanoma is a poorly immunogenic cell line derived from B78-H1 melanoma cells, which were originally derived from B16 melanoma.^51–53^ These cells were obtained from Ralph Reisfeld (Scripps Research Institute) in 2002. B78 cells were transfected with functional GD2/GD3 synthase to express the disialoganglioside GD2,^51,53^ which is overexpressed on the surface of many human tumors including melanoma.^54^ These B78 cells were also found to lack melanin. B78 cells were grown in RPMI-1640 (Gibco) supplemented with 10% FBS and 1% penicillin/streptomycin, with periodic supplementation with 400 μg G418 and 500 μg Hygromycin B per mL. Mycoplasma testing was performed every 6 months. B78 tumors were engrafted by intradermal flank injection of 2×10^6^ tumor cells.^15^ We have previously developed successful immunotherapy regimens for mice bearing these B78 tumors, enabling the cure of mice with measurable tumors (~100 mm^3^ volume).^13,14,16^ These cured mice have demonstrated tumor-specific T-cell mediated memory, as detected by rejection of rechallenge with the same vs. immunologically distinct tumors. Here, we continue our investigation of the B78 tumor model and expand our work by investigating collagen changes that occur during therapy. Tumor size was determined using calipers and volume approximated as:

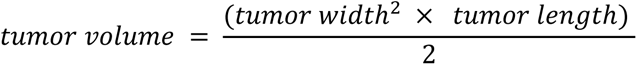

#### 2.1.2 Therapy Administration

Mice were randomized into treatment groups when tumors reached enrollment size (~150 mm^3^), which typically required 3-4 weeks of *in vivo* growth. The first day of treatment is defined here as ‘Day 0’ (**Figure 1a**). On Day 0, external beam radiation therapy was delivered to the tumor surface of treated mice only, using an X-RAD 320 cabinet irradiator system (Precision X-Ray, North Branford, CT). Mice were immobilized using custom lead jigs that exposed only the dorsal right flank. Radiation was delivered in one fraction to a maximum dose of 12 Gray (Gy). Systemic mouse α-CTLA-4 antibody was administered to treated mice once daily on Days 2, 5, and 8 via intraperitoneal injections of 200 μg in 200 μL phosphate buffered saline (PBS) (**Figure 1a**). The α-CTLA-4 antibody was provided by Bristol-Meyers Squibb (Redwood City, CA). Hu14.18-IL2 immunocytokine (IC), a monoclonal anti-GD2 antibody fused to IL2 cytokine, was administered to treated mice once daily on Days 5-9 via intratumoral injections of 50 μg in 100 μL PBS (**Figure 1a**). The Hu14.18-IL2 antibody was provided by AnYxis Immuno-Oncology GmbH (Vienna, Austria). Vehicle mice were injected once daily on Days 2, 5, and 8 via intraperitoneal injections of 200 μL PBS and once daily on Days 5-9 via intratumoral injections of 100 μL PBS. No external beam radiation therapy was administered to vehicle-treated mice. Pretreatment mice received no PBS or immunotherapy injections and no external beam radiation therapy.

**Figure 1:**
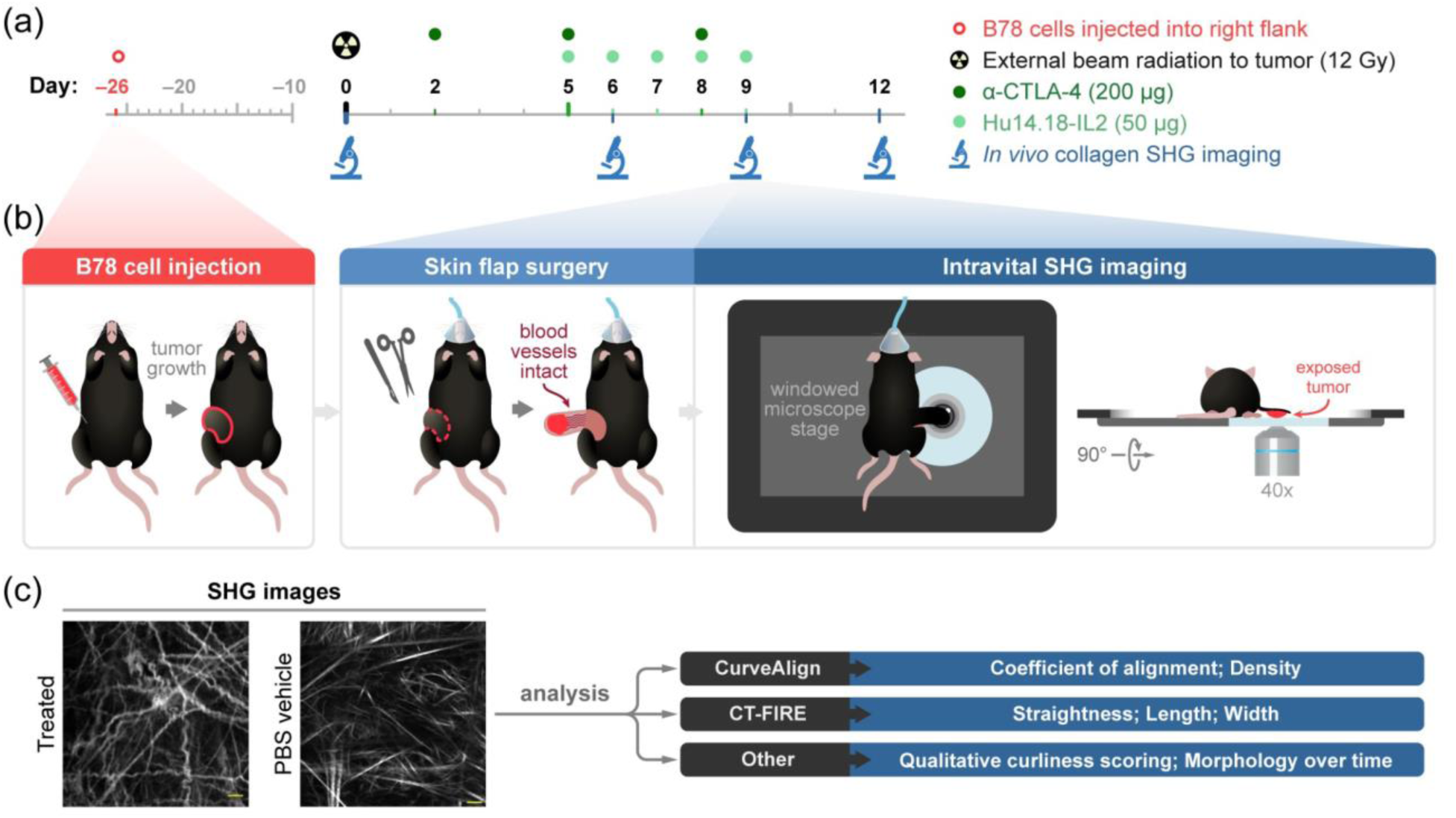
*In vivo* SHG imaging and treatment experimental workflow. **(a)** Experimental workflow began with intradermal inoculation of 2×10^6^ B78 melanoma cells into the right flank of our reporter mice. Tumors were monitored weekly until they reached ~150 mm^3^ volume (requiring ~26 days). For the treated group, combination therapy began on Day 0 with external beam radiation to the tumor surface (12 Gray) followed by intraperitoneal administration of α-CTLA-4 (200 μg) on Day 2, 5, 8 and intratumoral administration of Hu14.18-IL2 (50 μg) a monoclonal anti-GD2 antibody fused to IL2 on Days 5-9. For the vehicle group, matched PBS injections were administered at the same volumes and frequency as α-CTLA-4 and Hu14.18-IL2 in the treated mice and no radiation was administered. SHG imaging of tumor collagen was performed on pretreatment mice on Day 0 and both vehicle and treated mice on Days 6, 9, and 12. **(b)** B78 tumor growth was followed. On the indicated days of imaging, each mouse was anesthetized, and tumor skin flap surgery was performed where dermal and subcutaneous skin layers were gently cut away from the peritoneum revealing the tumor with intact vasculature. The tumor and skin flap were placed on a glass slide for SHG imaging and the mouse on a specially designed microscope stage. Throughout *in vivo* imaging, each mouse was kept under anesthesia and inside a heating chamber that enclosed the microscope stage. **(c)** The acquired SHG collagen images were then analyzed using CurveAlign and CT-FIRE software to quantify morphological and phenotypic changes. Qualitative analysis of the collagen images was also performed to evaluate curliness and morphology over time.

#### 2.1.3 Intravital Tumor Imaging

Intravital imaging of the mouse melanoma tumors was performed in pretreatment mice (n = 4), vehicle mice (n = 6), and treated mice (n = 6). Images were acquired on Days 0, 6, 9, and 12 of therapy (**Figure 1a**), in two independent experiments, to capture temporal collagen changes throughout the course of treatment. Day 0 images, pretreatment group, are used as a baseline comparison for both vehicle and immunotherapy treated groups. Immediately prior to tumor imaging, skin flap surgery exposed flank tumors. Mice were anesthetized with isoflurane, then the skin around the tumor was cut into a flap and separated from the body cavity so that the tumor laid flat on the imaging stage while still connected to the vasculature.^55–57^ Mice were placed on a specialized microscope stage for imaging and kept in a heating chamber (air maintained at 37 °C) during imaging. An imaging dish insert and PBS for coupling were used with surgical tape to secure skin flap tumors (**Figure 1b**).

### 2.2 Collagen SHG Imaging

Collagen SHG images were captured with a custom-built multi-photon microscope (Bruker) using an ultrafast femtosecond laser (InSight DSC, Spectra Physics) with linear polarization. Images of collagen fibers were detected using a bandpass filter of 514/30 nm with a 1041 nm excitation (typical power 2.1-3.4 mW). All images were acquired with a 40×/1.13 NA water-immersion objective (Nikon) at 512×512 pixel resolution, 4.8 μs pixel dwell time, 32 average frames, and an optical zoom of 1.0. SHG images were acquired to sample collagen changes during therapy across 2-7 fields of view and multiple depths within each tumor.

### 2.3 Collagen SHG Image Analysis

Several quantitative parameters were extracted using two packages of curvelet-based analysis software: CurveAlign and CT-FIRE (**Figure 1c**). CurveAlign was used here for bulk analysis of collagen at the FOV level, while CT-FIRE was used to quantify collagen at an individual collagen fiber level and at the FOV level.

#### 2.3.1 CurveAlign Analysis

CurveAlign^6,20,25–27^ was used to quantify collagen features at a FOV level, including collagen coefficient of alignment and density from the SHG images. 8-bit SHG collagen images were imported into CurveAlign, and the fraction of coefficients to keep was set to 0.04. CurveAlign was used to calculate the coefficient of alignment for each FOV on a scale between 0 and 1, where 0 is unaligned and 1 is fully aligned collagen fibers. Along with the coefficient of alignment, CurveAlign was also used to calculate the density of collagen within the FOV. Density calculations required a region of interest (ROI) analysis. For our purposes, we set the entire FOV to be the ROI. The threshold was set to 50 based on optimization calculations to remove as much background as possible, and the CurveAlign density parameter was calculated as the number of pixels within the FOV that were above the threshold. To calculate the true density, the CurveAlign density was divided by the total area of the FOV in pixels, resulting in a true density value between 0 and 1 where zero is no pixels within the FOV contain collagen fibers, and one is every pixel within the FOV contains collagen fibers. CurveAlign was used to analyze images at a FOV level for pretreatment mice (n = 4 mice, 18 FOV), vehicle-treated mice (n = 6 mice, 32 FOV), and immunotherapy-treated mice (n = 6 mice, 29 FOV).

### 2.3.1 CT-FIRE Analysis

The CT-FIRE^20,25,26^ module was designed to extract features such as collagen fiber straightness, length, and width at an individual collagen fiber level. Collagen straightness was defined here as the distance between the collagen fiber end points divided by the distance along the path of the fiber. A straightness value of 1 indicates a perfectly straight fiber, while a straightness value of 0 indicates a highly curly fiber. The length parameter measured the distance along the fiber from one end to the other while the width was the average width along the fiber. Both length and width were measured in pixels. CT-FIRE was used to analyze pretreatment mice single collagen fibers (n = 4 mice, 4282 single fibers), vehicle-treated mice single collagen fibers, (n = 6 mice, 7081 single fibers), and immunotherapy-treated mice single collagen fibers (n = 6 mice, 6755 single fibers) at the individual curvelet level.

### 2.4 Immunofluorescence

Excised mouse melanoma tumor tissues were formalin fixed and paraffin-embedded for antibody staining with a fluorescent marker of fibroblasts (α-smooth muscle actin α-SMA, abcam SP171 ab150301). Embedded sections were deparaffinized and hydrated prior to antigen retrieval and placement in blocking solution. Next, the primary antibody was applied upon removal of blocking solution at the following dilution and incubation time: α-SMA – 1:200 for 15 min at room temperature. A secondary rabbit antibody was then added following the primary antibody incubation at 1:500 for 10 min at room temperature. Then a staining dye was added after secondary antibody washes: α-SMA – Opal-dye 520 (Akoya Biosciences OP-001001) at 1:100. Finally, stained sections were incubated in DAPI for 5 min at room temperature for nuclear labeling and mounted on coverslips for imaging. Imaging was performed at 20× using a Vectra multispectral imaging system (Akoya Biosciences) and a spectral library was generated to separate spectral curves for each fluorophore. Resulting images were analyzed using Nuance and inForm software (Akoya Biosciences).

### 2.5 Statistical analysis

A one-way ANOVA with Tukey’s multiple comparisons test was performed to assess treatment group differences in collagen parameters between every combination of the three treatment groups: pretreatment, vehicle, and treated (**Figures 2–3, S1**). A log-transformation was first applied to the treatment group level data to avoid skewness. A one-way ANOVA with Šídák’s multiple comparisons test was performed to assess differences over time between vehicle and treated groups on each imaging day (**Figure 4–5, S2**). A log-transformation was first applied to the time-course data to avoid skewness. Collagen results are represented as box and whisker plots showing median ± min / max, with mean represented as a dot. Collagen curves are represented as mean ± standard deviation. A one-way ANOVA with Tukey’s multiple comparisons test was performed to assess differences in fibroblast numbers between treated mouse tumors over time (**Figure S3**). Fibroblast results are represented as scatter plots showing mean ± standard deviation (GraphPad Prism 9.5).

**Figure 2:**
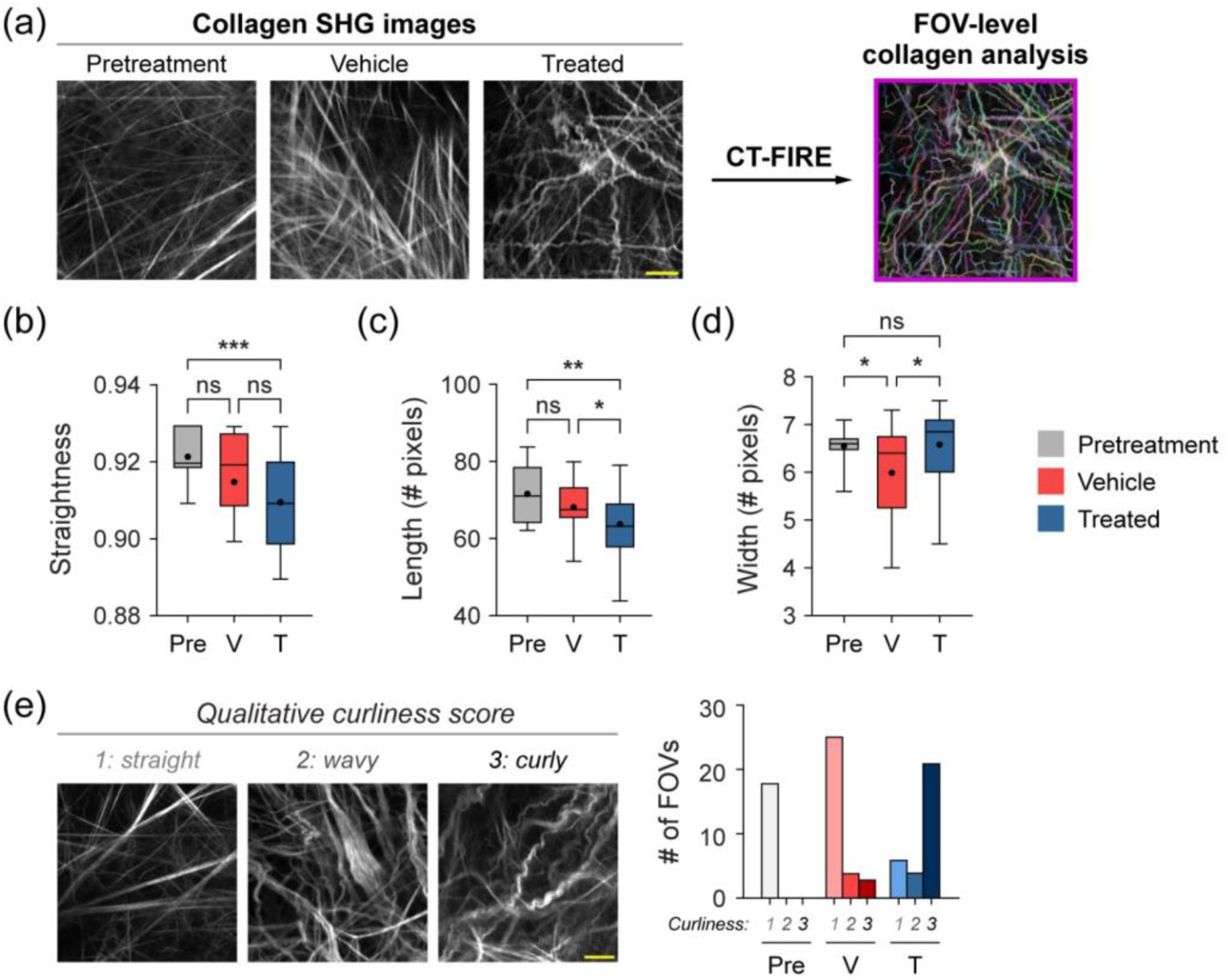
FOV-level analysis of collagen features by treatment group with CT-FIRE. **(a)** Representative *in vivo* SHG collagen images in B78 mouse melanoma tumors either from pretreatment, vehicle, or treated mice. FOV-level collagen analysis was performed using CT-FIRE. **(b)** FOV-level collagen straightness from pretreatment, vehicle, and treated mice (mean straightness: 0.92 pretreatment, 0.92 vehicle, 0.91 treated). A straightness value of 1 indicates a perfectly straight fiber, while a straightness value of 0 indicates a highly curly fiber. **(c)** FOV-level collagen length from pretreatment, vehicle, and treated mice (mean length: 72 pretreatment, 68 vehicle, 64 treated). **(d)** FOV-level collagen width from pretreatment, vehicle, and treated mice (mean width: 6.6 pretreatment, 6.0 vehicle, 6.6 treated). Box and whisker: median ± min/max, mean = dot. **(e)** Qualitative curliness scoring of SHG images from pretreatment, vehicle, and treated mice (n = 79 FOV) into three bins: 1 straight, 2 wavy, and 3 curly. Example images with their corresponding qualitative curliness score are shown. n = 4-6 mice per treatment group, pretreatment images n = 18 FOV, vehicle images n = 32 FOV, treated images n = 29 FOV. One-way ANOVA with Tukey’s multiple comparisons test, *p<0.05, **p<0.01, ***p<0.0005. Field of view, FOV. Scale bar 50 μm.

## 3 Results

### 3.1 FOV-level collagen analysis separates mice by treatment group

Mouse melanoma tumors were imaged *in vivo* in pretreatment mice (n = 4), vehicle-treated mice with no radiation (referred to as vehicle mice) (n = 6), and radiation plus immunotherapy-treated mice (referred to as treated mice) (n = 6). CT-FIRE was used to calculate curvelet based collagen changes at the FOV level from SHG images of all three treatment groups (**Figure 2a**). Representative *in vivo* SHG images show striking collagen fiber morphology differences across treatment groups (**Figure 2a**). Quantitatively, collagen from treated mice was significantly less straight than collagen from pretreatment mice (**Figure 2b**, ***p<0.001). Straightness differences between treated and vehicle mice were not statistically significant although treated mice appeared to have less straight collagen (**Figure 2b**). No significant difference in straightness was seen between vehicle and pretreatment mice either (**Figure 2b**). Second, collagen fibers from treated mice were significantly shorter in length compared to both pretreatment and vehicle mice (**Figure 2c**, treated vs. pretreatment **p<0.01, treated vs. vehicle *p<0.05) with no significant difference in length between vehicle and pretreatment. Third, collagen from treated mice was significantly wider than vehicle mice (**Figure 2d**, *p<0.05) though not significantly different compared to pretreatment mice. Collagen from vehicle mice is also thinner compared to pretreatment mice (**Figure 2d**, *p<0.05). Finally, qualitative curliness scoring of collagen images at the FOV level was performed by scoring of blinded images into bins of 1, 2, or 3 based on whether the collagen fibers were very straight (score 1), wavy (score 2), or very curly (score 3). Representative images for each score are shown in **Figure 2e**. Qualitative curliness scoring illustrated that pretreatment mice images were all scored at 1, indicating homogenous, very straight collagen across all FOV (**Figure 2e**). In contrast, vehicle mice showed some heterogeneity with these images scored mostly as 1 (n = 25 FOV) with a few images scored as 2 (n = 4 FOV) or 3 (n = 3 FOV) (**Figure 2e**). Treated mice also showed heterogeneity with very few images scored as 1 (n = 6 FOV) or 2 (n = 4 FOV) and most images scored as 3 (n = 21 FOV), indicating very curly collagen (**Figure 2e**). Overall, qualitative curliness scoring resulted in mostly straight collagen in pretreatment and vehicle mice with mostly curly collagen in treated mice (**Figure 2e**). Conversely, FOV level changes in collagen calculated with CurveAlign (**Figure S1a**) showed no significant difference in collagen coefficient of alignment between treatment groups (**Figure S1b**). Similarly, CurveAlign found no significant differences in collagen density across the three treatment groups at a FOV level, although treated mice appeared to have the highest density (**Figure S1c**).

### 3.2 CT-FIRE assesses changes in single fiber collagen morphology and separates mice by treatment group

Quantitative curvelet based collagen analysis was performed with CT-FIRE to assess changes at the single fiber level (**Figure 3a**), in contrast to the FOV-level analysis described in section 3.1. Single collagen fibers from treated mice were significantly less straight than collagen from vehicle and pretreatment mice (**Figure 3b**, treated vs. vehicle ***p<0.001, treated vs. pretreatment ****p<0.0001) Single collagen fibers from vehicle mice were also significantly less straight than collagen from pretreatment mice (**Figure 3b**, ****p<0.0001). Similarly, single collagen fibers from treated mice were significantly shorter than collagen from vehicle and pretreatment mice (**Figure 3c**, treated vs. vehicle ****p<0.0001, treated vs. pretreatment ****p<0.0001). Single fibers from vehicle mice were significantly shorter compared to pretreatment mice (**Figure 3c**, *p<0.05). Single collagen fibers from treated mice were also significantly wider than both vehicle and pretreatment mice (**Figure 3d**, treated vs. vehicle ****p<0.0001, treated vs. pretreatment ****p<0.001). Single collagen fibers from vehicle mice were significantly less wide compared to pretreatment mice (**Figure 3d**, ****p<0.0001). Overall, treated mice had collagen fibers that were curlier, shorter in length, and wider compared to both vehicle and pretreatment mice.

**Figure 3:**
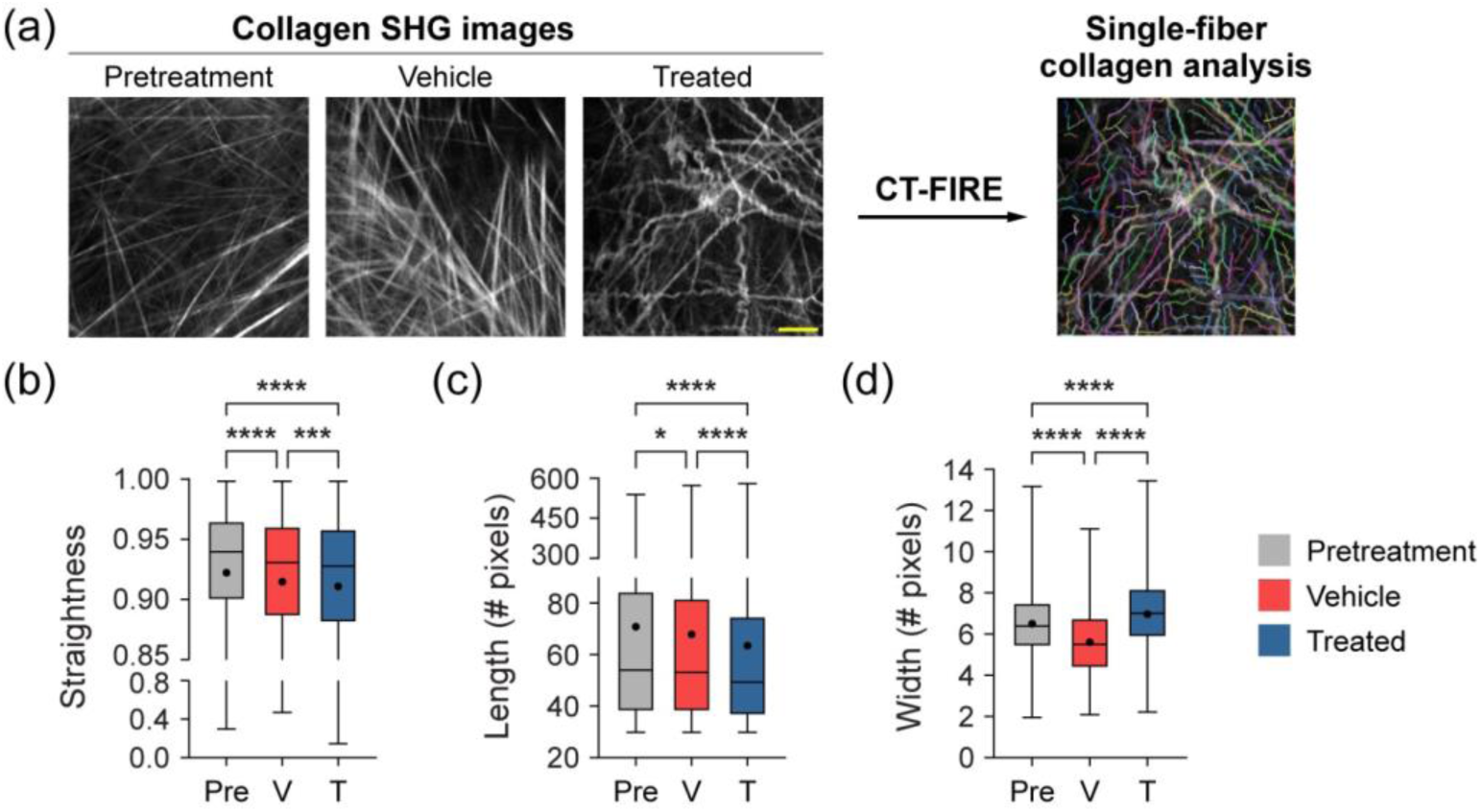
Single-fiber level analysis of collagen features by treatment group with CT-FIRE. **(a)** Representative *in vivo* SHG collagen images in B78 mouse melanoma tumors either from pretreatment, vehicle, or treated mice. Single fiber collagen analysis was performed using CT-FIRE. **(b)** Single fiber level collagen straightness from pretreatment, vehicle, and treated mice (mean straightness: 0.92 pretreatment, 0.92 vehicle, 0.91 treated). A straightness value of 1 indicates a perfectly straight fiber, while a straightness value of 0 indicates a highly curly fiber. **(c)** Single fiber level collagen length from pretreatment, vehicle, and treated mice (mean length: 71 pretreatment, 68 vehicle, 64 treated). **(d)** Single fiber level collagen width from pretreatment, vehicle, and treated mice (mean width: 6.5 pretreatment, 5.6 vehicle, 7.0 treated). Box and whisker: median ± min/max, mean = dot. n = 4-6 mice per treatment group, pretreatment curvelets n = 4282, vehicle curvelets n = 9229, treated curvelets n = 6755. One-way ANOVA with Tukey’s multiple comparisons test *p<0.05, ***p<0.0005, ****p<0.0001. Scale bar is 50 μm.

### 3.3 CT-FIRE and CurveAlign show time-dependent collagen morphology changes at the FOV level

FOV-level analysis for each day of treatment was performed using CT-FIRE and CurveAlign to extract time and treatment dependent changes. Representative *in vivo* SHG images show striking collagen fiber morphology differences across treatment groups and time-course (**Figure 4a**). With CT-FIRE analysis, vehicle mouse melanoma tumors, PBS injections only with no radiation, showed aligned, straight collagen fibers at Day 6, 9, and 12 (**Figure 4a**) that mirror the Day 0 pretreatment phenotype. In contrast, treated mouse melanoma tumors, radiation and immunotherapy treated, showed curly, tortuous collagen fibers at Day 6, 9, and especially 12 (**Figure 4a**). Surprisingly, no significant changes in collagen straightness were seen across the treatment course, at the FOV level, when comparing treated and vehicle mice (**Figure 4b**). Treated mice appeared to have less straight, curlier collagen compared to vehicle mice on Days 9 and 12, although these differences were insignificant. Treated mice showed significantly shorter collagen fiber length compared to vehicle mice at Day 12 only (**Figure 4c**, *p<0.05). Finally, treated mice exhibited significantly wider collagen fibers compared to vehicle mice at Day 6 only (**Figure 4d**). Collagen changes within a treatment group over time were also observed at the FOV level. Within the treated group, collagen fiber length shortened significantly from Day 9 to Day 12 (**Figure 4c**, *p<0.05) with no differences in straightness or width (**Figure 4b, 4d**). Within the vehicle group, collagen fiber width increased significantly between Day 6 to Day 9 (**Figure 4d**, *p<0.05) with no differences in straightness or length (**Figure 4b, 4c**). With CurveAlign (**Figure S2a**), no differences in collagen coefficient of alignment were observed over the treatment course, at the FOV level, when comparing treated and vehicle mice (**Figure S2b**). In contrast, treated mice did exhibit higher collagen density at Day 6 of treatment when compared to vehicle mice (**Figure S2c**, *p<0.05). Density differences between treated and vehicle mice were not different on any other day (**Figure S2c**). Within the vehicle group over time, collagen coefficient of alignment changed significantly from Day 6 to Day 9 (**Figure S2b**, *p<0.05) with no changes in density (**Figure S2c**). No differences were observed over time within the treated group for coefficient of alignment or density at the FOV level (**Figure S2b-c**).

**Figure 4:**
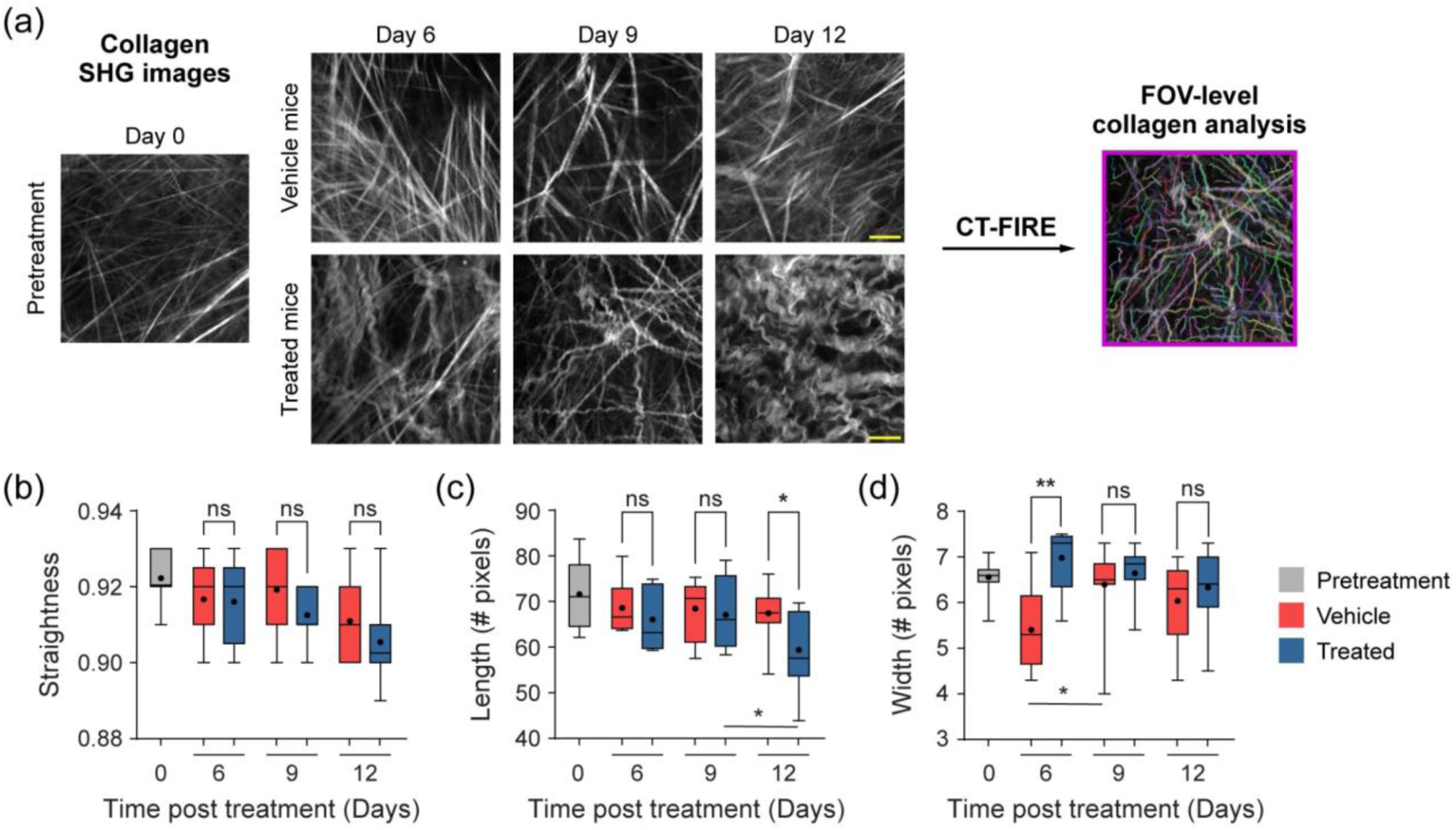
FOV-level analysis of collagen features over time with treatment using CT-FIRE. **(a)** Representative *in vivo* SHG collagen images in B78 mouse melanoma tumors from Day 0 pretreatment mice and Day 6, 9, 12 vehicle and treated mice. FOV level collagen analysis was performed using CT-FIRE. **(b)** FOV-level collagen straightness from pretreatment, vehicle, and treated mice over time with treatment (mean straightness vehicle:treated Day 6 0.92:0.92, Day 9 0.92:0.91, Day 12 0.91:0.91). A straightness value of 1 indicates a perfectly straight fiber, while a straightness value of 0 indicates a highly curly fiber. **(c)** FOV-level collagen length from pretreatment, vehicle, and treated mice over time with treatment (mean length vehicle:treated Day 6 69:66, Day 9 68:67, Day 12 67:59, mean length treated:treated Day 9 to Day 12 67:59). **(d)** FOV-level collagen width from pretreatment, vehicle, and treated mice over time with treatment (mean width vehicle:treated Day 6 5.4:7.0, Day 9 6.4:6.6, Day 12 6.0:6.3, mean width vehicle:vehicle Day 6 to Day 9 5.4:6.4). Note, the Day 9 vehicle median value is present but very close to the minimum value. Box and whisker: median ± min/max, mean = dot. n = 4-6 mice per treatment group, pretreatment images n = 18 FOV, vehicle images n = 32 FOV, treated images n = 29 FOV. One-way ANOVA with Šídák’s multiple comparisons test *p<0.05, **p<0.005. Field of view, FOV. Scale bar is 50 μm.

### 3.4 CT-FIRE shows time-dependent changes in single fiber collagen morphology

Finally, single-fiber level analysis was performed for each day of treatment using CT-FIRE to extract time and treatment dependent changes (**Figure 5a**). Treated mice exhibited less straight collagen fibers compared to vehicle mice on Day 9 (***p<0.001) and Day 12 (*p<0.05) but not Day 6 (**Figure 5b**). Treated mice also exhibited shorter collagen fibers on Day 9 (*p<0.05) and Day 12 (****p<0.0001) but not on Day 6 (**Figure 5c**). Similarly, the number of αSMA+ cells, presumed to be mostly fibroblasts, decreased over time in treated mouse melanoma tumors, with a significant decrease on Day 9 of treatment compared to Day 6 (**Figure S3**). Interestingly, collagen fibers from treated mice were significantly wider compared to vehicle mice on all days of treatment (**Figure 5d**, Day 6 ****p<0.0001, Day 9 *p<0.05, Day 12 ****p<0.0001). Single fiber collagen changes were also observed within each treatment group over time. Within the vehicle group, collagen fiber straightness changed significantly between Day 9 and Day 12 (**Figure 5b**, ***p<0.001) and collagen width changed significantly between all three days (**Figure 5d**, Day 6 and 9 ****p<0.0001, Day 9 and 12 ****p<0.0001, Day 6 and 12 ****p<0.001) with no differences in length (**Figure 5c**). Within the treated group, collagen fiber straightness, length, and width all changed significantly between Day 6 and D12 as well as Day 9 and Day 12 (**Figure 5b-d**, straightness Day 6 and 12 *p<0.05, Day 9 and 12 **p<0.01, length Day 6 and 12 ***p<0.001, Day 9 and 12 ****p<0.001, width Day 6 and 12 ****p<0.001, Day 9 and 12 ****p<0.0001) with only width changing significantly between Day 6 and Day 9 (**Figure 5d**, ****p<0.001). Qualitative scoring of collagen images shows that pretreatment mouse tumors were scored exclusively as 1, indicating a straight phenotype (**Figure 5e-5f**). Similarly, 78% of vehicle mouse tumor images were scored as 1, a straight phenotype, across all treatment days (**Figure 5e**). As a result, very few vehicle mouse tumor images were scored as 2 (n = 4 FOV) or 3 (n = 3 FOV) across treatment time (**Figure 5e**). In contrast, treated mouse tumors were qualitatively scored mostly as 3 (68%), a curly phenotype, across all treatment days (**Figure 5f**). Only a few images (n = 4 FOV) were scored as 2, wavy, or (n = 6 FOV) 1, straight (**Figure 5f**). Interestingly, the few treated mouse images that were scored as 1, straight, were only from Day 6 and Day 9 post-treatment, with all collagen fibers scored as wavy or curly (2 or 3) by Day 12 (**Figure 5f**).

**Figure 5:**
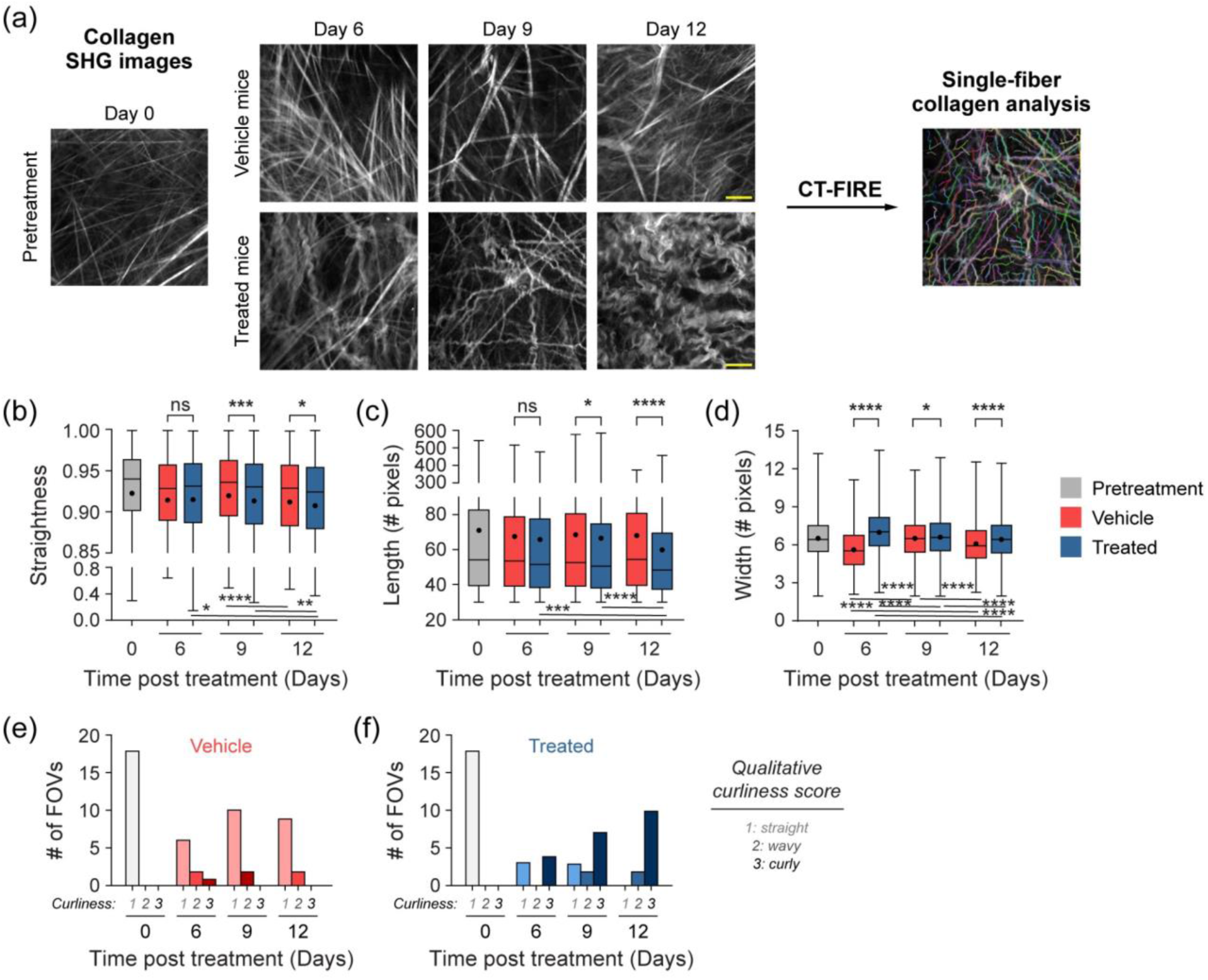
Single-fiber level analysis of collagen features over time with treatment using CT-FIRE. **(a)** Representative *in vivo* SHG collagen images in B78 mouse melanoma tumors from Day 0 pretreatment mice and Day 6, 9, 12 vehicle and treated mice. Single-fiber analysis was performed using CT-FIRE. **(b)** Single-fiber collagen straightness from pretreatment, vehicle, and treated mice over time with treatment (mean straightness vehicle:treated Day 6 0.91:0.91, Day 9 0.92:0.91, Day 12 0.91:0.91, mean straightness vehicle:vehicle Day 9 to Day 12 0.92:0.91, mean straightness treated: treated Day 6 to Day 12 0.91:0.91, Day 9 to Day 12 0.91:0.91). A straightness value of 1 indicates a perfectly straight fiber, while a straightness value of 0 indicates a highly curly fiber. **(c)** Single-fiber collagen length from pretreatment, vehicle, and treated mice over time with treatment (mean length vehicle:treated Day 6 67:66, Day 9 68:67, Day 12 68:60, mean length treated:treated Day 6 to Day 12 66:67, Day 9 to Day 12 67:60). **(d)** Single-fiber collagen width from pretreatment, vehicle, and treated mice over time with treatment (mean width vehicle:treated Day 6 5.6:7.0, Day 9 6.5:6.7, Day 12 6.1:6.5, mean width vehicle:vehicle Day 6 to Day 9 5.6:6.5, Day 9 to Day 12 6.5:6.1, Day 6 to Day 12 5.6:6.1, mean width treated:treated Day 6 to Day 9 7.0:6.7, Day 9 to Day 12 6.7:6.5, Day 6 to Day 12 7.0:6.5). Box and whisker: median ± min/max, mean = dot. **(e-f)** Qualitative curliness scoring of SHG images from pretreatment, vehicle, and treated mice over time with treatment (n = 79 FOV) into three bins: 1 straight, 2 wavy, and 3 curly. n = 4-6 mice per treatment group, pretreatment curvelets n = 4282, vehicle curvelets n = 9229, treated curvelets n = 6755. One-way ANOVA with Šídák’s multiple comparisons test *p<0.05, **p<0.01, ***p<0.001, ****p<0.0001. Scale bar is 50 μm.

## 4 Discussion

During tumorigenesis, cancer cells and cancer associated fibroblasts secrete excess metalloproteinases and collagen, which often leads to the reconstruction, linearization, and remodeling of collagen in the tumor microenvironment.^3^ This remodeling may be especially relevant for melanoma patients whose skin is 70% collagen and whose tumors are often fibroblast rich.^29–32,58^ Though SHG imaging has enabled insights into collagen trends in breast and pancreatic cancer, few studies have performed SHG of collagen in melanoma.^22,49,50^ To our knowledge, this is the first study to quantify *in vivo* mouse melanoma tumor collagen reorganization during immunotherapy treatment using label-free SHG imaging. We quantified six collagen morphology features during combination radiotherapy and immunotherapy or PBS vehicle using CurveAlign and CT-FIRE software. We examined collagen reorganization and phenotypic changes with temporal context at both the FOV-level and the single-fiber level. We found that collagen length and width changes were the most sensitive parameters to distinguish treated and control tumors. We also demonstrated that single-fiber level collagen analysis via CT-FIRE was most sensitive to the changes between treatment groups. Temporally, these collagen changes occurred primarily on Day 9 and 12 of treatment.

*In vivo* SHG imaging provided clear visualization of collagen fibers within mouse tumors. We observed distinct collagen fiber morphology that varied by treatment group and time. Our imaging captured collagen reorganization that occurred only in the radiation and immunotherapy treated mice at Day 9 and 12 of treatment. Overall, we showed that live SHG imaging of mouse melanoma tumors produced high contrast images where mice could be categorized by treatment group based on collagen phenotype that changed during therapy.

We reported FOV-level collagen changes using both CurveAlign and CT-FIRE. Our FOV-level collagen coefficient of alignment measurements performed in CurveAlign were not statistically different across all three groups. As coefficient of alignment is dependent on both the straightness of collagen as well as the fiber orientations in relation to each other, this calculation is most accurate when a defined tumor border can be marked within the FOV.^25^ Our *in vivo* images were acquired within the tumor so no such border was present, which may partly explain the limited effectiveness of this parameter. If desired, we could acquire *in vivo* mouse SHG images at the tumor edge to draw a border within each FOV, which should improve the relevance of the coefficient of alignment. Interestingly, our FOV-level density measurements showed that treated mouse tumors appeared to have higher collagen density, but this was only statistically significant on Day 6. Though increased collagen density leads to negative outcomes in many cancers, similar findings have been shown in human melanoma and human / rat breast cancer where increased collagen density was associated with better tumor outcomes.^6,50,59^ This increased collagen density in the skin may help prevent tumor cell metastasis, linking it to a healthy phenotype.^2^ In contrast, CT-FIRE showed statistically significant differences in length and width of collagen fibers from treated mice versus PBS control. At the FOV level, collagen fibers from treated mice versus control mice are shorter in length. This indicates that collagen fibers from treated mice are less linearized and may be transitioning towards a healthier phenotype. This shorter collagen characteristic may also indicate cleavage of collagen fibers by MMPs from local fibroblasts.^22^ Additionally, collagen fibers from treated mice versus PBS control mice are wider, which further supports our assessment that treated mouse tumors contain collagen that is less linearized. Perhaps this increase in width is a result of collagen fibers that are less stretched and less taut. Others have similarly shown that shorter, wider, curlier collagen is a characteristic of healthy human skin compared to basal cell carcinoma biopsies.^60,61^

We also showed single fiber collagen changes using CT-FIRE. Radiation and immunotherapy treated mouse tumors possessed collagen fibers that were significantly less straight, shorter, and wider compared to PBS control tumors. This treated tumor collagen phenotype is consistent with what is expected for healthy tissue collagen versus tumor associated collagen signatures (TACS) where straightness and alignment are increased.^5,20^ The decrease in straightness or alignment in our treated mouse tumors was also consistent with *in vivo* collagen SHG of mouse healthy ears (less aligned) compared to mouse ears bearing B16-F10 melanoma (more aligned).^49^ Additionally, shorter and wider collagen fibers restrict tumor cell invasion, via decreased β1 integrins and metalloproteinase associated genes as well as increased E-cadherin on tumor cells, in human and rat breast cancers.^59^ The difference in collagen phenotype between our PBS vehicle tumors and treated mouse tumors may be due to the treatment itself or changes occurring during tumor regression.

Collagen changes from treated mouse tumors versus vehicle tumors also varied with time. Differences in single fiber collagen straightness and length were not statistically significant until Days 9 and 12 of treatment. Interestingly, single fiber collagen width changed the fastest, with the largest differences observed on Day 6 of treatment. This may suggest that collagen straightness and length alterations are more linked to tumor regression or changes in tumor cell signaling, as they changed most at the end of treatment when tumors shrink in response to therapy.^30^ These late collagen straightness and length changes may also reflect immune infiltrate changes in the tumor as treated mice have received the full regimen of α-CTLA-4 and immunocytokine by Day 9, activating T and NK cells while reducing T regulatory cells.^13,14^ In contrast, perhaps collagen width alterations that are most different at Day 6 of treatment are more linked to the radiation therapy that occurred on Day 0. Radiation therapy can affect fibroblasts within the tissue, leading to overactivation and sometimes increased collagen synthesis.^62^ We showed that treated tumor αSMA+ fibroblast numbers increased significantly from Day 0 to Day 6 and collagen density significantly increased at Day 6, possibly reflecting the overactivation effect of radiation therapy on the fibroblasts. However, we also showed a significant decrease in treated tumor αSMA+ fibroblast numbers from Day 6 to Day 9. This Day 9 reduction in αSMA+ fibroblasts may be partly why we see collagen fiber straightness and length change on Day 9 and 12, as well as indicate reduced collagen deposition^63^ near the end of treatment, though additional assays are needed to confirm this. In a cohort of male head and neck cancer patients, 20 Gy of radiation therapy led to a decrease in collagen III synthesis.^62^ Interestingly, in a cohort of female breast cancer patients who received 50 Gy, the opposite effect was seen following radiation therapy where an increase in collagen I and III synthesis was observed.^64^ We presume the collagen imaged within these melanoma tumors is primarily collagen I with some collagen III contribution.^65^ Additional studies are needed to determine whether collagen synthesis is increased or decreased in our melanoma model; however, our results may follow the trend of the head and neck cancer patients due to tissue similarity, where radiation therapy reduced collagen synthesis. The collagen morphology changes we observed may also be linked to the reprogramming of cancer associated macrophages and fibroblasts, which have been implicated in collagen deposition and pro-tumor behavior.^20,37,66–70^ Others have shown collagen SHG morphology changes with chemotherapy and targeted therapy, but to our knowledge, this is the first study to quantify these changes during combination radiation and immunotherapy *in vivo*.^38^ As immunotherapy is increasingly common for melanoma and other cancers, it may be clinically important to monitor other biomarkers of response.^12,71^ This work suggests that collagen morphology changes may be prognostic during immunotherapy.

Overall, we found that CT-FIRE was more sensitive to collagen changes with this model and treatment, at both the FOV-level and single-fiber level, compared to CurveAlign. This may be because CT-FIRE considers single collagen fibers through curvelet calculations compared to CurveAlign that calculates collagen changes at a FOV-level only. This was especially true as our dataset did not contain tumor borders that are needed for relevant coefficient of alignment calculations.

It is important to note that although nearly all of the SHG signal we gathered is expected to be collagen, there is a chance that other ECM proteins were captured such as elastin.^17,72^ We do not expect that the small contribution of elastin to our SHG signal significantly impacts our conclusions.

Although *in vivo* SHG imaging provides information on collagen dynamics across treatment groups and treatment time-courses, SHG imaging alone is not sufficient to specify a biological mechanism for collagen changes. Parallel measurements such as flow cytometry, western blot, and single cell RNA sequencing, some of which we are currently pursuing, should identify specific cells, cytokines, and proteins that drive this reorganization. We are also interested in which component or components of the triple therapy prompted these collagen changes. Future studies may include similar collagen SHG analysis in mice that received only a single component of the therapeutic regimen, as well as expansion of this work to a mouse colon carcinoma model for comparison. Ultimately, this imaging approach could provide insight into tumor microenvironmental changes and collagen dynamics, with key implications in improving immunotherapy response in cancer. This work could also lead to exploration of collagen biomarkers for additional cancer types.

## 5 Conclusion

Here, collagen morphology features were quantified from *in vivo* SHG images of mouse melanoma tumors during radiation and immunotherapy. Collagen dynamics and phenotypic changes with temporal context were quantitatively examined with CurveAlign and CT-FIRE software to evaluate collagen changes at the FOV level and the single fiber level. Collagen from radiation and immunotherapy treated mice reorganized during treatment toward a healthier tissue phenotype including: shorter, wider, curlier collagen fibers, with modestly higher collagen density. Temporally, collagen fiber straightness and length changed late in treatment (Day 9 and Day 12) while width and density changed early in treatment (Day 6) compared to vehicle mice. Overall, we have shown substantial quantitative and qualitative changes in collagen during the melanoma response to this radio-immunotherapy regimen. SHG imaging in preclinical models or patient samples may provide insight into tumor microenvironmental features that are associated with improved immunotherapy response in cancer, and new biomarkers of response.

## Acknowledgments

This work was supported by a Morgridge Interdisciplinary Postdoctoral Fellowship to A.R.H; The Midwest Athletes against Childhood Cancer (MACC) Fund; Stand Up to Cancer, The St. Baldrick’s Foundation; The Crawdaddy Foundation; The Cancer Research Institute; The Alex’s Lemonade Stand Foundation; The Carol Skornicka Chair in Biomedical Imaging; Retina Research Foundation Daniel M. Albert Chair; and the National Institutes of Health [R35CA197078, 5P30CA014520-40, UL1TR002373, P01CA250972, R01CA205101, U54CA232568, R01CA278051, R01CA272855]. The authors would like to acknowledge the University of Wisconsin Carbone Cancer Center Support Grant P30CA014520 and the University of Wisconsin Small Animal Imaging & Radiotherapy Facility. The authors would like to acknowledge Karla Esbona, Rebecca Baus, and the University of Wisconsin Translational Research Initiatives in Pathology laboratory (TRIP), supported by the UW Department of Pathology and Laboratory Medicine, UWCCC (P30 CA014520) and the Office of The Director – NIH (S10OD023526) for use of its facilities and services. The authors would like to thank Yuming Liu for helpful guidance, Amy Erbe-Gurel, Alexander Rakhmilevich, and Anne-Sophie Mancha for helpful discussions, as well as Matt Stefely for figure formatting assistance.

## Code, Data, and Materials Availability

The raw data supporting the conclusions of this article will be made available by the authors to any qualified researcher. The download and use of CurveAlign and CT-FIRE is free and open to the public at the following GitHub repository: https://github.com/uw-loci/curvelets/releases/tag/5.0

## Ethics Statement

The animal study was reviewed and approved by the Institutional Animal Care and Use Committee at University of Wisconsin-Madison.

## References

1. N. I. Nissen, M. Karsdal, and N. Willumsen, “Collagens and Cancer associated fibroblasts in the reactive stroma and its relation to Cancer biology,” Journal of Experimental & Clinical Cancer Research 38(1), 115 (2019) [doi:10.1186/s13046-019-1110-6].

2. C. Bonnans, J. Chou, and Z. Werb, “Remodelling the extracellular matrix in development and disease,” 12, Nat Rev Mol Cell Biol 15(12), 786–801, Nature Publishing Group (2014) [doi:10.1038/nrm3904].

3. L. E. Iannucci et al., “Optical Imaging of Dynamic Collagen Processes in Health and Disease,” Frontiers in Mechanical Engineering 8 (2022).

4. B. Erdogan et al., “Cancer-associated fibroblasts promote directional cancer cell migration by aligning fibronectin,” J Cell Biol 216(11), 3799–3816 (2017) [doi:10.1083/jcb.201704053].

5. P. P. Provenzano et al., “Collagen reorganization at the tumor-stromal interface facilitates local invasion,” BMC Medicine 4(1), 38 (2006) [doi:10.1186/1741-7015-4-38].

6. J. S. Bredfeldt et al., “Automated quantification of aligned collagen for human breast carcinoma prognosis,” J Pathol Inform 5(1), 28 (2014) [doi:10.4103/2153-3539.139707].

7. H. Salmon et al., “Matrix architecture defines the preferential localization and migration of T cells into the stroma of human lung tumors,” J Clin Invest 122(3), 899–910 (2012) [doi:10.1172/JCI45817].

8. M. Arnold et al., “Global Burden of Cutaneous Melanoma in 2020 and Projections to 2040,” JAMA Dermatology 158(5), 495–503 (2022) [doi:10.1001/jamadermatol.2022.0160].

9. R. L. Siegel et al., “Cancer statistics, 2023,” CA A Cancer J Clinicians 73(1), 17–48 (2023) [doi:10.3322/caac.21763].

10. N. Zaidi and E. M. Jaffee, “Immunotherapy transforms cancer treatment,” Journal of Clinical Investigation 129(1), 46–47 (2018) [doi:10.1172/JCI126046].

11. J. D. Wolchok et al., “Long-Term Outcomes With Nivolumab Plus Ipilimumab or Nivolumab Alone Versus Ipilimumab in Patients With Advanced Melanoma,” Journal of Clinical Oncology, Wolters Kluwer Health (2021) [doi:10.1200/JCO.21.02229].

12. M. R. Albertini, “The age of enlightenment in melanoma immunotherapy,” j. immunotherapy cancer 6(1), 80, s40425-018-0397–0398 (2018) [doi:10.1186/s40425-018-0397-8].

13. Z. S. Morris et al., “*In Situ* Tumor Vaccination by Combining Local Radiation and Tumor-Specific Antibody or Immunocytokine Treatments,” Cancer Research 76(13), 3929– 3941 (2016) [doi:10.1158/0008-5472.CAN-15-2644].

14. Z. S. Morris et al., “Tumor-Specific Inhibition of *In Situ* Vaccination by Distant Untreated Tumor Sites,” Cancer Immunology Research 6(7), 825–834 (2018) [doi:10.1158/2326-6066.CIR-17-0353].

15. P. M. Carlson et al., “Depth of tumor implantation affects response to in situ vaccination in a syngeneic murine melanoma model,” J Immunother Cancer 9(4), e002107 (2021) [doi:10.1136/jitc-2020-002107].

16. T. J. Aiken et al., “Short-course neoadjuvant in situ vaccination for murine melanoma,” J Immunother Cancer 10(1), e003586 (2022) [doi:10.1136/jitc-2021-003586].

17. W. R. Zipfel et al., “Live tissue intrinsic emission microscopy using multiphoton-excited native fluorescence and second harmonic generation,” Proceedings of the National Academy of Sciences 100(12), 7075–7080, Proceedings of the National Academy of Sciences (2003) [doi:10.1073/pnas.0832308100].

18. R. Gauderon, P. B. Lukins, and C. J. R. Sheppard, “Optimization of second-harmonic generation microscopy,” Micron 32(7), 691–700 (2001) [doi:10.1016/S0968-4328(00)00066-4].

19. G. Cox et al., “3-Dimensional imaging of collagen using second harmonic generation,” Journal of Structural Biology 141(1), 53–62 (2003) [doi:10.1016/S1047-8477(02)00576-2].

20. J. S. Bredfeldt et al., “Computational segmentation of collagen fibers from second-harmonic generation images of breast cancer,” J Biomed Opt 19(1), 16007 (2014) [doi:10.1117/1.JBO.19.1.016007].

21. M. W. Conklin et al., “Aligned Collagen Is a Prognostic Signature for Survival in Human Breast Carcinoma,” The American Journal of Pathology 178(3), 1221–1232 (2011) [doi:10.1016/j.ajpath.2010.11.076].

22. E. Brown et al., “Dynamic imaging of collagen and its modulation in tumors in vivo using second-harmonic generation,” 6, Nat Med 9(6), 796–800, Nature Publishing Group (2003) [doi:10.1038/nm879].

23. R. M. Williams, W. R. Zipfel, and W. W. Webb, “Interpreting Second-Harmonic Generation Images of Collagen I Fibrils,” Biophysical Journal 88(2), 1377–1386 (2005) [doi:10.1529/biophysj.104.047308].

24. K. M. Riching et al., “3D collagen alignment limits protrusions to enhance breast cancer cell persistence,” Biophys J 107(11), 2546–2558 (2014) [doi:10.1016/j.bpj.2014.10.035].

25. Y. Liu et al., “Methods for Quantifying Fibrillar Collagen Alignment,” Methods Mol Biol 1627, 429–451 (2017) [doi:10.1007/978-1-4939-7113-8_28].

26. Y. Liu and K. W. Eliceiri, “Quantifying Fibrillar Collagen Organization with Curvelet Transform-Based Tools,” J Vis Exp (165) (2020) [doi:10.3791/61931].

27. F. S. P. Campagnola Paul J., Ed., “Quantitative Approaches for Studying the Role of Collagen in Breast Cancer Invasion and Progression,” in Second Harmonic Generation Imaging, CRC Press (2013).

28. A. Oikarinen, “Aging of the skin connective tissue: how to measure the biochemical and mechanical properties of aging dermis,” Photodermatol Photoimmunol Photomed 10(2), 47– 52 (1994).

29. L. Ziani et al., “Melanoma-associated fibroblasts decrease tumor cell susceptibility to NK cell-mediated killing through matrix-metalloproteinases secretion,” Oncotarget 8(12), 19780–19794 (2017) [doi:10.18632/oncotarget.15540].

30. J. Mazurkiewicz et al., “Melanoma cells with diverse invasive potential differentially induce the activation of normal human fibroblasts,” Cell Commun Signal 20(1), 63 (2022) [doi:10.1186/s12964-022-00871-x].

31. L. Van Hove and E. Hoste, “Activation of Fibroblasts in Skin Cancer,” Journal of Investigative Dermatology 142(4), 1026–1031 (2022) [doi:10.1016/j.jid.2021.09.010].

32. F. Papaccio et al., “Profiling Cancer-Associated Fibroblasts in Melanoma,” Int J Mol Sci 22(14), 7255 (2021) [doi:10.3390/ijms22147255].

33. T. Liu et al., “Cancer-associated fibroblasts: an emerging target of anti-cancer immunotherapy,” J Hematol Oncol 12(1), 86 (2019) [doi:10.1186/s13045-019-0770-1].

34. L. Monteran and N. Erez, “The Dark Side of Fibroblasts: Cancer-Associated Fibroblasts as Mediators of Immunosuppression in the Tumor Microenvironment,” Frontiers in Immunology 10 (2019).

35. P. P. Provenzano et al., “Collagen density promotes mammary tumor initiation and progression,” BMC Med 6, 11 (2008) [doi:10.1186/1741-7015-6-11].

36. D. Kedrin et al., “Intravital imaging of metastatic behavior through a Mammary Imaging Window,” Nat Methods 5(12), 1019–1021 (2008) [doi:10.1038/nmeth.1269].

37. O. Maller et al., “Tumour-associated macrophages drive stromal cell-dependent collagen crosslinking and stiffening to promote breast cancer aggression,” Nat. Mater. 20(4), 548–559 (2021) [doi:10.1038/s41563-020-00849-5].

38. A. J. Walsh et al., “Collagen density and alignment in responsive and resistant trastuzumab-treated breast cancer xenografts,” J. Biomed. Opt 20(2), 026004 (2015) [doi:10.1117/1.JBO.20.2.026004].

39. M. Papanicolaou et al., “Temporal profiling of the breast tumour microenvironment reveals collagen XII as a driver of metastasis,” Nat Commun 13(1), 4587 (2022) [doi:10.1038/s41467-022-32255-7].

40. J. J. Northey et al., “Stiff stroma increases breast cancer risk by inducing the oncogene ZNF217,” Journal of Clinical Investigation 130(11), 5721–5737 (2020) [doi:10.1172/JCI129249].

41. J. M. Szulczewski et al., “Directional cues in the tumor microenvironment due to cell contraction against aligned collagen fibers,” Acta Biomaterialia 129, 96–109 (2021) [doi:10.1016/j.actbio.2021.04.053].

42. K. Esbona et al., “The Presence of Cyclooxygenase 2, Tumor-Associated Macrophages, and Collagen Alignment as Prognostic Markers for Invasive Breast Carcinoma Patients,” The American Journal of Pathology 188(3), 559–573 (2018) [doi:10.1016/j.ajpath.2017.10.025].

43. H. Laklai et al., “Genotype tunes pancreatic ductal adenocarcinoma tissue tension to induce matricellular fibrosis and tumor progression,” Nat Med 22(5), 497–505 (2016) [doi:10.1038/nm.4082].

44. N. S. Nagathihalli et al., “Signal Transducer and Activator of Transcription 3, Mediated Remodeling of the Tumor Microenvironment Results in Enhanced Tumor Drug Delivery in a Mouse Model of Pancreatic Cancer,” Gastroenterology 149(7), 1932–1943.e9 (2015) [doi:10.1053/j.gastro.2015.07.058].

45. D. Tokarz et al., “Characterization of Pancreatic Cancer Tissue Using Multiphoton Excitation Fluorescence and Polarization-Sensitive Harmonic Generation Microscopy,” Front Oncol 9, 272 (2019) [doi:10.3389/fonc.2019.00272].

46. W. Hu et al., “Nonlinear Optical Microscopy for Histology of Fresh Normal and Cancerous Pancreatic Tissues,” PLOS ONE 7(5), e37962, Public Library of Science (2012) [doi:10.1371/journal.pone.0037962].

47. C. R. Drifka et al., “Periductal stromal collagen topology of pancreatic ductal adenocarcinoma differs from that of normal and chronic pancreatitis,” Modern Pathology 28(11), 1470–1480 (2015) [doi:10.1038/modpathol.2015.97].

48. C. R. Drifka et al., “Highly aligned stromal collagen is a negative prognostic factor following pancreatic ductal adenocarcinoma resection,” Oncotarget 7(46), 76197–76213 (2016) [doi:10.18632/oncotarget.12772].

49. P.-C. Wu et al., “In vivo Quantification of the Structural Changes of Collagens in a Melanoma Microenvironment with Second and Third Harmonic Generation Microscopy,” 1, Sci Rep 5(1), 8879, Nature Publishing Group (2015) [doi:10.1038/srep08879].

50. C. Thrasivoulou et al., “Optical delineation of human malignant melanoma using second harmonic imaging of collagen,” Biomed. Opt. Express 2(5), 1282 (2011) [doi:10.1364/BOE.2.001282].

51. M. Haraguchi et al., “Isolation of GD3 synthase gene by expression cloning of GM3 a-2,8-sialyltransferase cDNA using anti-GD2 monoclonal antibody,” Proc. Natl. Acad. Sci. USA 91(22), 10455–10459, (1994) [doi:10.1073/pnas.91.22.10455].

52. S. Silagi, “CONTROL OF PIGMENT PRODUCTION IN MOUSE MELANOMA CELLS IN VITRO,” Journal of Cell Biology 43(2), 263–274 (1969) [doi:10.1083/jcb.43.2.263].

53. J. C. Becker et al., “An antibody-interleukin 2 fusion protein overcomes tumor heterogeneity by induction of a cellular immune response.,” Proc. Natl. Acad. Sci. U.S.A. 93(15), 7826–7831 (1996) [doi:10.1073/pnas.93.15.7826].

54. B. Nazha, C. Inal, and T. K. Owonikoko, “Disialoganglioside GD2 Expression in Solid Tumors and Role as a Target for Cancer Therapy,” Front. Oncol. 10, 1000 (2020) [doi:10.3389/fonc.2020.01000].

55. J. R. W. Conway, S. C. Warren, and P. Timpson, “Context-dependent intravital imaging of therapeutic response using intramolecular FRET biosensors,” Methods 128, 78–94 (2017) [doi:10.1016/j.ymeth.2017.04.014].

56. A. L. B. Seynhaeve and T. L. M. ten Hagen, “Intravital Microscopy of Tumor-associated Vasculature Using Advanced Dorsal Skinfold Window Chambers on Transgenic Fluorescent Mice,” JoVE (131), 55115 (2018) [doi:10.3791/55115].

57. C. L. G. J. Scheele et al., “Multiphoton intravital microscopy of rodents,” Nature Reviews Methods Primers 2(89) (2022) [10.1038/s43586-022-00168-w].

58. “Dermis - an overview | ScienceDirect Topics,” <https://www.sciencedirect.com/topics/medicine-and-dentistry/dermis> (accessed 19 April 2023).

59. O. Maller et al., “Collagen architecture in pregnancy-induced protection from breast cancer,” Journal of Cell Science, jcs.121590 (2013) [doi:10.1242/jcs.121590].

60. M. Sendín-Martín, et al., “Quantitative collagen analysis using second harmonic generation images for the detection of basal cell carcinoma with ex vivo multiphoton microscopy,” Experimental Dermatology 32(4), 392–402 (2023) [doi:10.1111/exd.14713].

61. N. Kiss et al., “Quantitative Analysis on Ex Vivo Nonlinear Microscopy Images of Basal Cell Carcinoma Samples in Comparison to Healthy Skin,” Pathol. Oncol. Res. 25(3), 1015– 1021 (2019) [doi:10.1007/s12253-018-0445-1].

62. K. Mazurek et al., “Collagen Type III Metabolism Evaluation in Patients with Malignant Head and Neck Cancer Treated with Radiotherapy,” Biomed Res Int 2018, 8702605 (2018) [doi:10.1155/2018/8702605].

63. Z. Miskolczi et al., “Collagen abundance controls melanoma phenotypes through lineage-specific microenvironment sensing,” Oncogene 37(23), 3166–3182 (2018) [doi:10.1038/s41388-018-0209-0].

64. R. Keskikuru et al., “Radiation-induced changes in skin type I and III collagen synthesis during and after conventionally fractionated radiotherapy,” Radiotherapy and Oncology 70(3), 243–248, Elsevier (2004) [doi:10.1016/j.radonc.2003.11.014].

65. D. Singh, V. Rai, and D. K. Agrawal, “Regulation of Collagen I and Collagen III in Tissue Injury and Regeneration,” Cardiology and cardiovascular medicine 7(1), 5, NIH Public Access (2023) [doi:10.26502/fccm.92920302].

66. C. J. Hanley et al., “A subset of myofibroblastic cancer-associated fibroblasts regulate collagen fiber elongation, which is prognostic in multiple cancers,” Oncotarget 7(5), 6159– 6174 (2016) [doi:10.18632/oncotarget.6740].

67. L. A. Jolly et al., “Fibroblast-Mediated Collagen Remodeling Within the Tumor Microenvironment Facilitates Progression of Thyroid Cancers Driven by BrafV600E and Pten Loss,” Cancer Research 76(7), 1804–1813 (2016) [doi:10.1158/0008-5472.CAN-15-2351].

68. M. Schnoor et al., “Production of Type VI Collagen by Human Macrophages: A New Dimension in Macrophage Functional Heterogeneity,” The Journal of Immunology 180(8), 5707–5719 (2008) [doi:10.4049/jimmunol.180.8.5707].

69. R. Chen, L. Huang, and K. Hu, “Natural products remodel cancer-associated fibroblasts in desmoplastic tumors,” Acta Pharmaceutica Sinica B 10(11), 2140–2155 (2020) [doi:10.1016/j.apsb.2020.04.005].

70. J. N. Ouellette et al., “Navigating the Collagen Jungle: The Biomedical Potential of Fiber Organization in Cancer,” Bioengineering 8(2), 17 (2021) [doi:10.3390/bioengineering8020017].

71. N. A. Nixon et al., “Current landscape of immunotherapy in the treatment of solid tumours, with future opportunities and challenges,” Curr Oncol 25(5), e373–e384 (2018) [doi:10.3747/co.25.3840].

72. R. Datta et al., “Fluorescence lifetime imaging microscopy: fundamentals and advances in instrumentation, analysis, and applications,” J. Biomed. Opt. 25(07), 1 (2020) [doi:10.1117/1.JBO.25.7.071203].

