## Supplemental Figures S1-S3 for "Quantifying *in vivo* collagen reorganization during immunotherapy in murine melanoma with second harmonic generation imaging"

### Supplementary Material

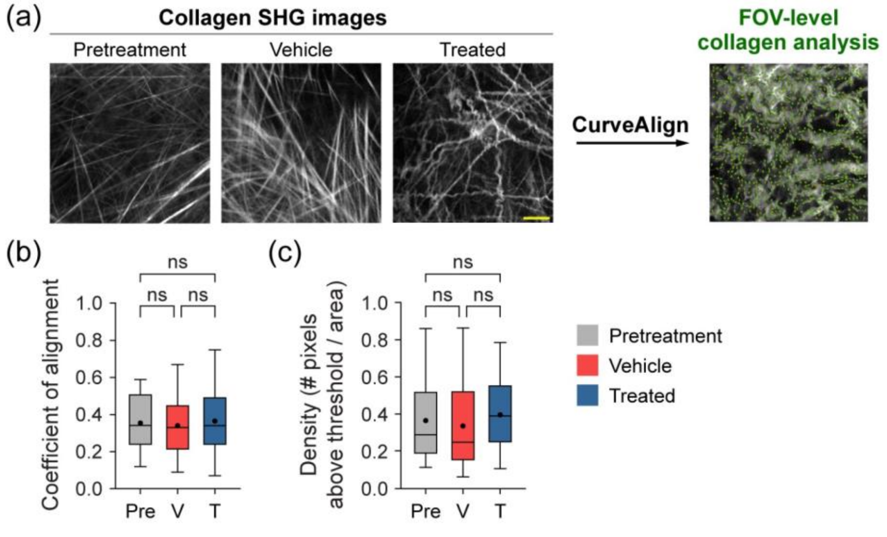


**Figure S1: FOV-level analysis of collagen coefficient of alignment and density changes by treatment group using CurveAlign. (a)** Representative *in vivo* SHG collagen images in B78 mouse melanoma tumors either from pretreatment, vehicle, or treated mice. FOV-level collagen analysis was performed using CurveAlign. **(b)** FOV-level collagen coefficient of alignment from pretreatment, vehicle, and treated mice (mean coefficient of alignment: 0.35 pretreatment, 0.34 vehicle, 0.37 treated). For coefficient of alignment, 0 indicates fibers are unaligned and 1 indicates fibers are fully aligned. **(c)** FOV-level collagen density from pretreatment, vehicle, and treated mice (mean density: 0.37 pretreatment, 0.34 vehicle, 0.40 treated). Box and whisker: median ± min/max, mean = dot. n = 4-6 mice per treatment group, pretreatment images n = 18 FOV, vehicle images n = 32 FOV, treated images n = 29 FOV. One-way ANOVA with Tukey's multiple comparison tests. Field of view, FOV. Scale bar is 50 μm.

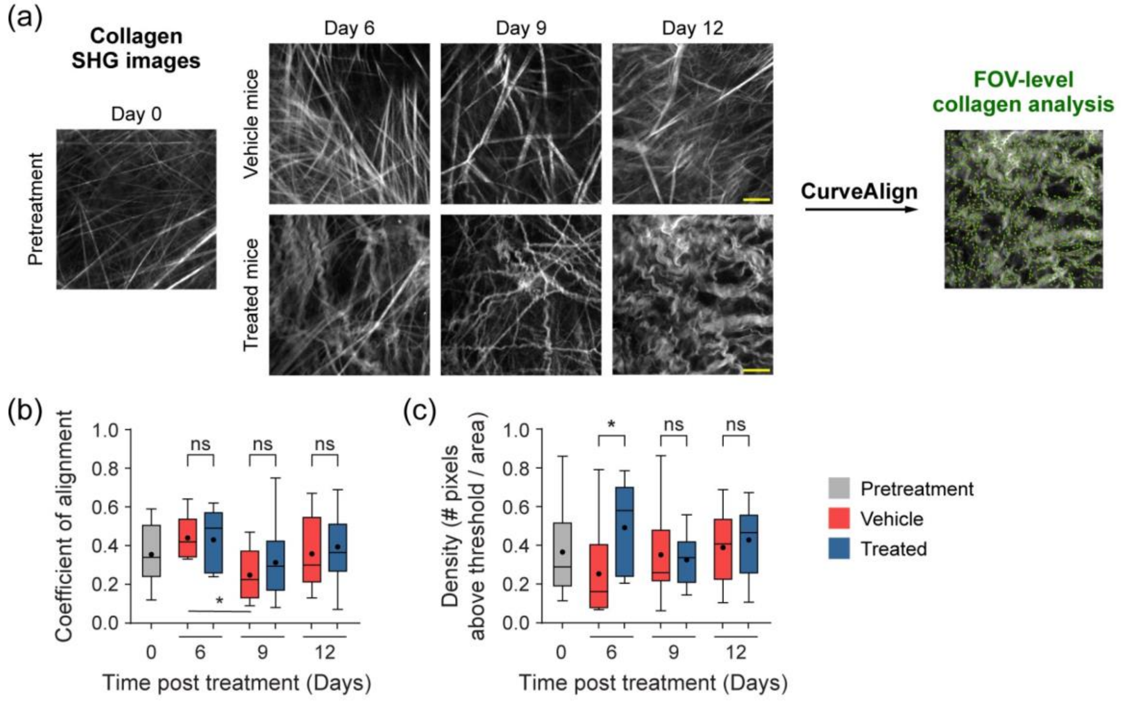


**Figure S2: FOV-level analysis of collagen coefficient of alignment and density changes over time with treatment using CurveAlign. (a)** Representative *in vivo* SHG collagen images in B78 mouse melanoma tumors from Day 0 pretreatment mice and Day 6, 9, 12 vehicle and treated mice. FOV-level collagen analysis was performed using CurveAlign. **(b)** FOV-level collagen coefficient of alignment from pretreatment, vehicle, and treated mice over time with treatment (mean coefficient of alignment vehicle:treated Day 6 0.44:0.43, Day 9 0.25:0.31, Day 12 0.36:0.39, mean coefficient of alignment vehicle:vehicle Day 6 to Day 9 0.44:0.25). For coefficient of alignment, 0 indicates fibers are unaligned and 1 indicates fibers are fully aligned. **(c)** FOV-level collagen density from pretreatment, vehicle, and treated mice over time with treatment (mean density vehicle:treated Day 6 0.25:0.49, Day 9 0.35:0.33, Day 12 0.39:0.43). Box and whisker: median ± min/max, mean = dot. n = 4-6 mice per treatment group, pretreatment images n = 18 FOV, vehicle images n = 32 FOV, treated images n = 29 FOV. One-way ANOVA with Šídák's multiple comparison tests, *$p<0.05$. Field of view, FOV. Scale bar is 50 μm.

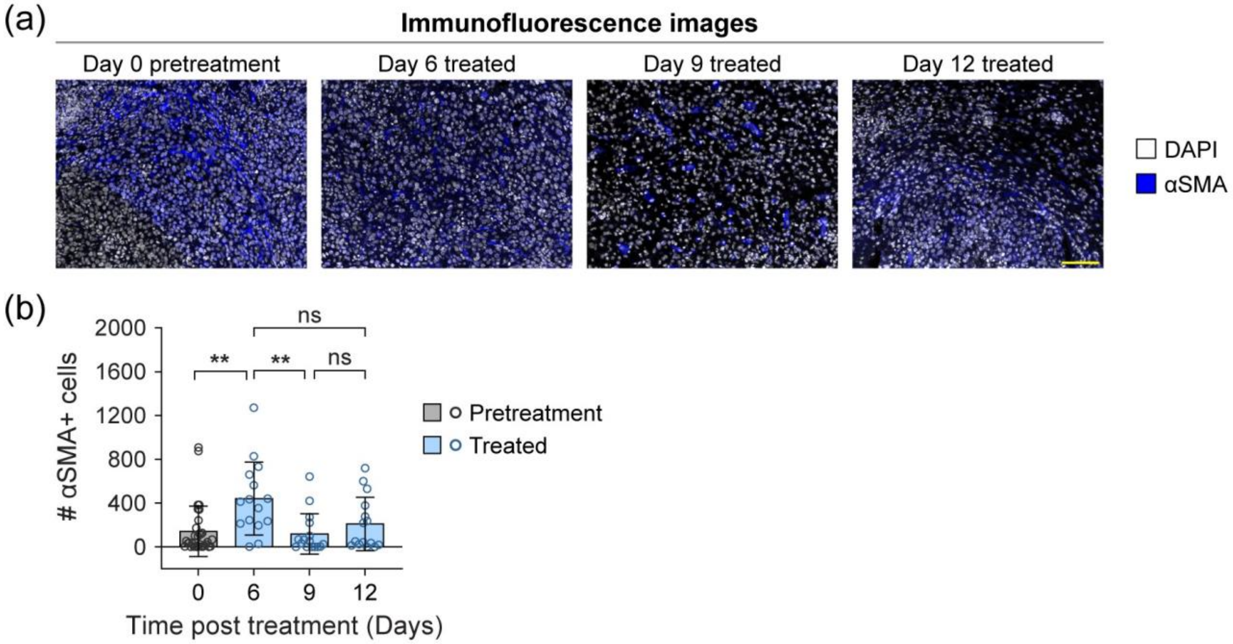


**Figure S3: Immunofluorescence of αSMA+ fibroblast populations in mouse melanoma tumors during treatment. (a)** Representative 20× immunofluorescence images in B78 mouse melanoma tumors from Day 0 pretreatment mice and Day 6, 9, 12 treated mice (DAPI gray, αSMA+ fibroblasts blue). **(b)** Number of fibroblasts within melanoma tumors from pretreatment and treated mice over time (mean number of fibroblasts Day 0: 143, Day 6: 441, Day 9: 119, Day 12: 210). Bars: mean ± SD, each point represents a FOV. n = 2-4 mice per treatment group, pretreatment Day 0 images n = 32 FOV, treated Day 6 images n = 15 FOV, treated Day 9 images = 16 FOV, treated Day 12 images = 15 FOV. One-way ANOVA with Tukey's multiple comparison tests, **p<0.01. Field of view, FOV. Scale bar is 100 μm.
